# Genome-wide association studies reveal novel loci controlling tuber flesh color and oxidative browning in *Dioscorea alata*

**DOI:** 10.1101/2023.03.12.532275

**Authors:** Komivi Dossa, Angélique Morel, Mahugnon Ezékiel Houngbo, Ana Zotta Mota, Erick Malédon, Jean-Luc Irep, Jean-Louis Diman, Pierre Mournet, Sandrine Causse, Kien Nguyen Van, Denis Cornet, Hâna Chair

**Affiliations:** CIRAD, UMR AGAP Institut, 97170 Petit Bourg, Guadeloupe, France; UMR AGAP Institut, Univ Montpellier, CIRAD, INRAE, Institut Agro, F-34398 Montpellier, France; CIRAD, UMR AGAP Institut, F-34398 Montpellier, France; UR1321 ASTRO Agrosystèmes tropicaux, INRAE, Petit□Bourg (Guadeloupe), F□97170, France; Plant Resources Center, An Khanh, Hoai Duc, Hanoi, Vietnam

**Keywords:** Yam, association genetics, quality traits, molecular markers, breeding

## Abstract

**Background:** Consumers’ preferences for food crops are guided by quality attributes. This study aimed at deciphering the genetic basis of quality traits, especially tuber flesh color (FC) and oxidative browning (OB) in *Dioscorea alata*, based on the genome-wide association studies (GWAS) approach. The *D. alata* panel was planted at two locations in Guadeloupe. At harvest, the FC was scored visually as white, cream, or purple on longitudinally sliced mature tubers. The OB was scored visually as the presence or absence of browning after 15 minutes of exposure of the sliced samples to ambient air.

**Results:** Phenotypic characterization for FC and OB of a diverse panel of *D. alata* genotypes highlighted significant variation within the panel and across two locations. The genotypes within the panel displayed a weak structure and could be classified into 3 subpopulations. GWAS identified 14 and 4 significant associations for tuber FC and OB, respectively, with phenotypic variance, explained values ranging from 7.18 to 18.04%. Allele segregation analysis at the significantly associated loci highlighted the favorable alleles for the desired traits, i.e., white FC and no OB. A total of 24 putative candidate genes were identified around the significant signals. A comparative analysis with previously reported quantitative trait loci indicated that numerous genomic regions control these traits in *D. alata*.

**Conclusion:** Our study provides important insights into the genetic control of tuber FC and OB in *D. alata*. The major and stable loci can be further utilized to improve selection in breeding programs for developing new cultivars with enhanced tuber quality.

## Introduction

*Dioscorea alata* L., commonly known as greater yam or water yam, contributes significantly to food security in Africa, the Caribbean islands, and Asia.^1,2^ Bred cultivars, with improved yields and disease resistance, have contributed substantially to meet market demands.^3-5^ Although *D. alata* is widely adapted to different environments and shows high yield and tolerance to biotic and abiotic stresses, the quality of its tubers is lower compared to other species, such as *D. rotoundata*.^6^ Improving quality attributes of yam tubers has been largely neglected. Therefore, satisfactory end-use quality remains the primary objective of yam breeding, along with the increased yield.^7^

Desirable aesthetic characteristics in food depend mainly on texture and color. A recent survey by Effah-Manu et al., emphasized tuber flesh color (FC) as a key quality trait in yams according to end-users preferences.^8^ The tuber FC range in greater yam varies from white, yellowish, orange, pink, and purple.^9^ However, color choice is linked to consumer preferences and the dish. For instance, the preparation of pounded yam (called “fufu” in West Africa) requires a white to yellowish color.^10^ In some regions, Reunion Island, Guyana, Philippine, and India, purple FC is preferred for the preparation of cakes and mousselines.^11^ Moreover, color variation is generally attributed to differential metabolic profiles, including flavonoids, anthocyanins, and carotenoids.^12,13^ Therefore, flesh-color variation and associated metabolites could substantially enhance the nutritional value of yam.

In yams, quality generally refers to the nutritional contents, physical appearance, and processing ease.^14^ Oxidative browning (OB) of tubers causes undesirable color changes and can cause bitterness in pre-processed yams with an undesirable flavor.^15,10,16^ Moreover, OB may prevent the preparation of local dishes such as pounded yam.^1^ Although phenotypic variation concerning OB has been widely studied, the genetic factors underlying this trait have not been explored fully.^17^ Rinaldo et al., showed that polyphenolic oxidation is a major cause of OB in *D. alata*.^18^ Deciphering the genetic mechanisms underlying OB is a critical strategy to avoid discoloration in yam tubers and meet the quality requirements of consumers.

Genetic gain estimates showed that conventional breeding practices alone are insufficient to satisfy market demands.^19^ Utilizing innovative technologies with conventional breeding can accelerate genetic gain.^20^ Progress in genotyping techniques and the development of modern genotype-phenotype association statistical models have provided a sophisticated way forward for identifying genomic regions and molecular markers associated with phenotypic variation in crops.^21^ These tools can be harnessed for trait introgression and rapid screening of progenies in breeding programs.^21^ Genome-wide association studies (GWAS) have been widely adopted in plant species to uncover marker-trait associations by providing an effective way to complement the traditional quantitative trait loci (QTL) mapping.^1,21^ Gatarira et al., used GWAS to identify six putative candidate genes on chromosome 5 associated with OB in *D. alata*.^1^ Their diversity panel was limited to West Africa, hence not harboring the wealth of alleles controlling this trait. Ehounou et al. identified five QTLs for OB and three for tuber FC in *D. alata* using the linkage mapping approach in bi-parental populations.^22^ However, the parents used in their study did not greatly vary for tuber FC, and their phenotyping procedures did not consider the environmental influence. It is well known that environmental conditions affect plant performance, yield, and quality, with significant implications in breeding programs.^2^ Hence, detecting stable QTLs not influenced by environmental factors is crucial for wide-scale use in breeding programs.

The present study aimed to decipher the genetic architectures of tuber FC and OB by using a diversified panel of *D. alata* for both traits.

## Materials and Methods

### Plant materials, growth conditions, and phenotyping

This study included 53 *D. alata* genotypes and one *D. cayenensis* clone used as an outgroup to verify the genotyping quality. This panel comprises accessions from the yam belt countries of West Africa, the Caribbean, and the Pacific islands (Table S1).

The materials were planted at two locations, Roujol (16°10′ 56′′ N, 61° 35′ 24′′ W, 10 meters above sea level, m.a.s.l) and Godet (16°20′ N, 61°30′ 0.10 m.a.s.l.), in Guadeloupe. The average temperature and relative humidity at Godet and Roujol during the experiment were 27.1°C / 79.5% and 25.8°C / 87.7%, respectively. A total of 30 seedlings of each genotype were planted and split into three replicates (10 seedlings per replication). These plants were spread over three ridges (65 m long) spaced 30 cm apart within the ridges. Sugarcane straw was used as mulch over the entire plot to limit weeds at Godet, while paper mulch was used at Roujol. The plot was drip irrigated, and planting started in March 2021. Yams were planted from seeds from the harvest of year n-1 when 50% of the tubers of the genotype germinated in the storage shed. The seeds (∼100 g) were treated with black soap and alcohol, by immersing them for 30s in a mixture (10 L of water, 0.5 L of black soap, and 0.5 L of alcohol) and sown 24 hours later. The tubers were harvested seven to nine months later, corresponding to the commercial harvest period. After harvest, tubers were stored in a storage room at 27 ◦C for three weeks.

All the genotypes were characterized for phenotypic data concerning two quality traits, FC and OB. Three well-developed tubers per genotype were peeled and sliced longitudinally, and their FC was immediately scored visually as 1: white, 2: cream, or 3: purple.^22^ The sliced samples were kept in an air-conditioned room at 25°C and exposed to ambient air for 15 minutes. The OB trait was scored visually as 1: change in color/browning or 2: no change in color/no browning.^22^ The score data were used for association analysis.

### Genotyping and single nucleotide polymorphism filtering

Genomic DNA of the 54 genotypes was extracted from the leaves using a mixed alkyltrimethylammonium bromide buffer and the kit NucleoMag Plant (Macherey-Nagel, Germany). DNA samples were quantified using ThermoFisher Scientific fluorometry Qubit, and fragment length was assessed with the Agilent Tapestation system. Illumina Tru-SEQ DNA PCR free and TruSeq Nano DNA Preparation Kit have been applied to generate sequencing libraries. Paired high-throughput sequencing (2 × 150 bp) was performed on an Illumina NovaSeq 6000 instrument on the GeT-PlaGe platform (Toulouse, France).

The quality of the whole genome sequencing (WGS) raw reads were first checked using the FastQC package version 0.11.7,^23^ trimmed by removing adaptor sequences and low-quality sequences using Trimmomatic version 0.39^24^ with the following parameters: SLIDINGWINDOW, 6:15; TRAILING, 20; HEADCROP, 3; and MINLEN, 90. Clean reads were then aligned to the *D. alata* v.2 reference genome using BWA-MEM version 0.7.15^25^ with default parameters. The alignment quality was analyzed with Qualimap version 2.2^26^ with default parameters. The calling of single nucleotide polymorphism (SNP) for each genotype was performed using GATK 4.1.6.0 with the HaplotypeCaller method.^27^

A consolidated variant call format (VCF) file for all genotypes was produced by VCFtools version 0.1.16,^28^ containing about 17 million SNPs. All InDel-like variants were eliminated. Then, the VCF file was filtered using the following parameters: --max-missing 0.2 – minQ 30 --minDP 8 --maf 0.05 –allele 2. With filters applied to reduce the quantity and eliminate bad-quality SNPs, there were 1.9 million SNPs left. The SNP density was estimated using the “CMplot” package^29^ in R4.0.23.^30^

### Population structure, principal component analysis, linkage disequilibrium, and SNP annotation

A python centralized package named “RESEQ2” has been designed internally in our research group, which gathers several next-generation sequencing data analysis packages (such as PLINK 1.07,^31^ ADMIXTURE 1.23,^32^ VCFtools 0.1.16,^28^ GATK 4.1.6.0, ANNOVAR 2.4^33^). The phylogenetic relationships among the accessions were measured based on the genetic distance using the neighbor-joining method (with 1000 bootstrap replicates) in PLINK 1.07.^31^ The resulting phylogenetic tree was visualized in MEGA-11. The population structure (K from 1-10; 100 runs per K) was assessed with ADMIXTURE 1.23,^32^ and the accessions with the membership coefficients (> 60%) were assigned to a specific Cluster. The remaining accessions were assigned to the admixture Cluster. The principal component analysis was conducted with PLINK 1.07^31^ with default parameters and plotted in R4.0.23. The linkage disequilibrium (LD) coefficients (*r*^2^) between pairwise high-quality SNPs were calculated in PLINK 1.07^31^ with default parameters. The maximum distance between two SNPs was defined as 310 kb. The LD decay graphs were produced in R4.0.23. The annotation of the filtered VCF file was performed with ANNOVAR 2.4^33^ with default parameters. Based on the annotation of the *D. alata* genome V.2,^34^ SNPs were categorized as occurring in coding regions (further grouped into synonymous or nonsynonymous SNPs), splicing sites, untranslated regions, intronic regions, upstream and downstream regions, and intergenic regions.

### Genome-wide association studies

Genome-wide association studies (GWAS) were performed with all 1.9 M SNPs using the statistical model Fixed and random model circulating probability unification (FarmCPU), and Mixed Linear Model implemented in the GAPIT3 package^35^ under R4.0.23^30^. The FarmCPU algorithm allows an improvement in the statistical power, an increase in the calculation efficiency, and a reduction of false associations in the GWAS approach.^36^ The FarmCPU formula is :

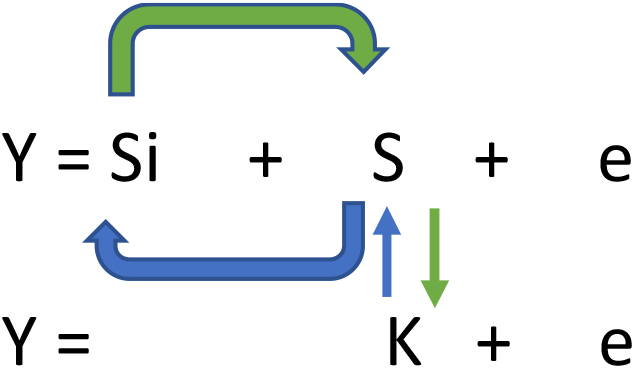

Where Y is the phenotype, Si is the testing marker, S is the pseudo-QTN, K is the kinship matrix, and e is residual. The kinship and first principal components are fitted as covariate variables in the FarmCPU to reduce the false positives. To eliminate the ambiguity of determining associated markers that are in LD with a testing marker, the model eliminates the confounding from kinship by using only the kinship derived from the associated markers.

The Manhattan plots were also generated in R4.0.23.^30^ SNPs having a significant association with traits were determined by the adjusted p-value. The threshold of P< 10^−8^ (0.05/n, with n = number of SNPs) was set to report a significant association. The quantile-quantile plots were generated by plotting the negative logarithms (−log_10_) *P*-values relative to their expected p-values to fit model relevance GWAS with the null hypothesis of no association and to determine to what extent the models considered the structure of the population. The effect of alleles at significant SNPs was assessed by comparing phenotyping data for allelic groups.

### Putative candidate gene identification

To inventory potential genes near associated SNP markers for target traits, we defined a window range of the average LD distance (5 kb upstream and 5 kb downstream) and used ANNOVAR 2.4^33^ annotated VCF file to locate genes around the significant SNPs.

### Validation of the significantly associated SNPs for tuber FC in an external panel

We obtained a phenotypic dataset on tuber FC of 30 *D. alata* accessions maintained and phenotyped in Vietnam. These accessions were planted at An Khánh District of Hoai Duc (Hanoï city) during the growing season 2016-2017. At harvest, FC of three mature tubers per accession was visually noted as explained above (1: white, 2: cream, or 3: purple).^22^ The accessions were genotyped (by WGS) as explained above, and a VCF file was generated. Peak SNPs initially identified through GWAS were searched in the VCF file of the Vietnam panel, and alleles at these SNPs were assessed by comparing tuber FC data for allelic groups.

### *In silico* comparative analysis of QTLs from previous studies on tuber FC and OB in *D. alata*

Quantitative trait loci (QTL) results from three previously published studies concerning QTLs related to tuber FC and OB in *D. alata* were extracted.^1,22,34^ Marker sequences for each reported QTL was retrieved, and local blast (Blastn) was used to identify corresponding physical positions in the reference genome V.2.^34^ The previously reported QTLs and those detected in our study were physically mapped using MapChart 2.32.^37^

## Results

### Phenotypic characterization in the *Dioscorea alata* diversity panel

The *D. alata* panel was classified into three categories based on the tuber FC: white, cream (or yellowish), and purple FCs (Figure 1A, 1B, 1C). Most accessions were white (a color generally preferred by consumers). There was variation within the two environments, suggesting that the environmental conditions influence the tuber FC in some genotypes. For instance, 46 and 68% of genotypes had white flesh tubers in Roujol and Godet, respectively (Figures 1D and 1E). Observed inconsistency of tuber FCs among planting sites was mainly from white to cream and vice versa.

**Figure 1.**
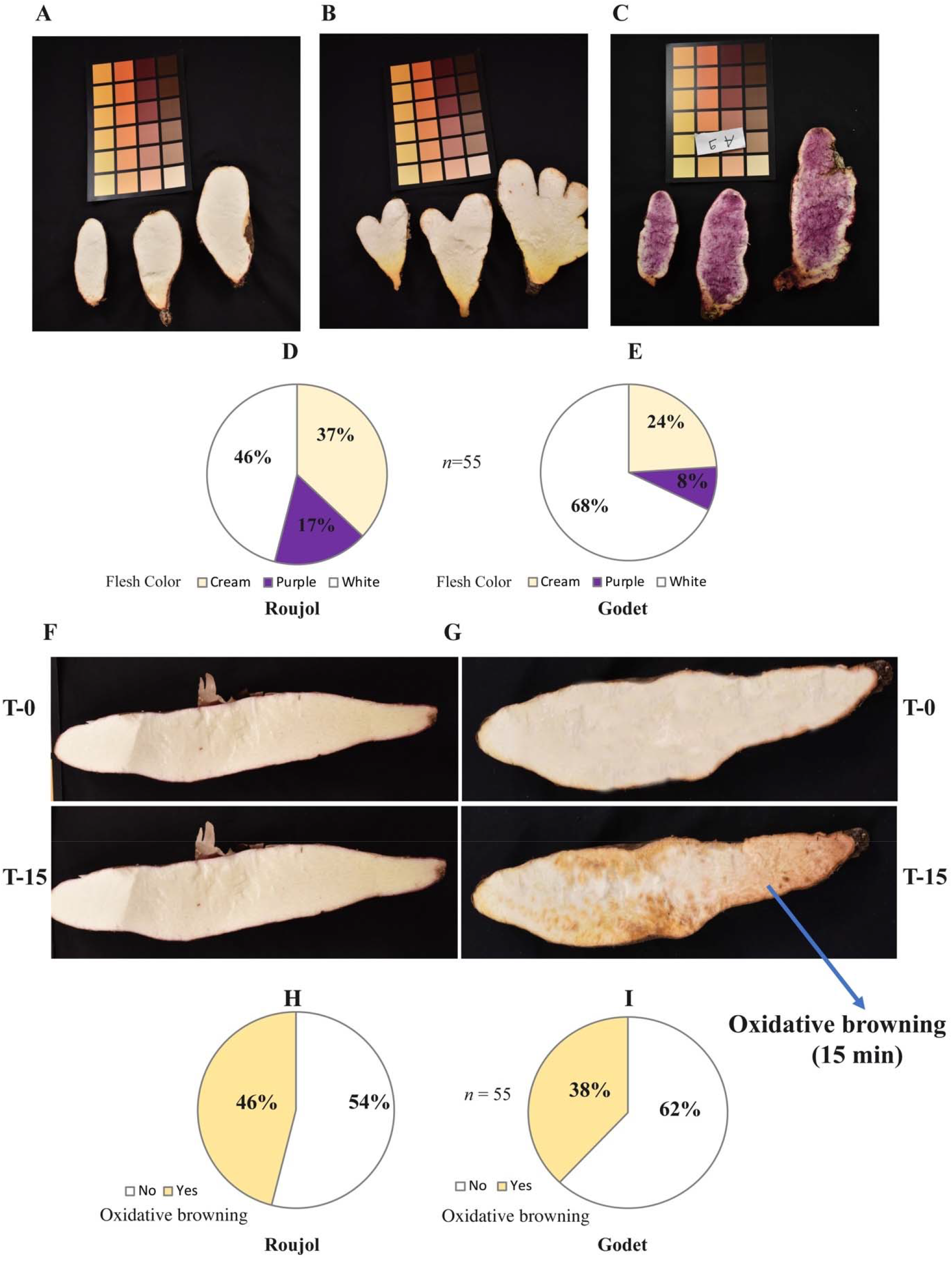
Tuber flesh color (FC) variation and oxidative browning (OB) in *Dioscorea alata* association panel. Examples of **A**) white colored, **B**) cream (yellow) colored, **C**) purple colored *tubers*; Variation in tuber FC within the panel at **D**) Roujol, **E**) Godet, Where “n” is the number of genotypes; **F**) Absence of OB; **G**) Presence of OB in a yam tuber after 15 minutes of cutting, where upper parts showed the color at T-0 (immediately after slicing), while lower parts show the color at T-15 (15 minutes after slicing); Variation of OB within the panel at **H**) Roujol and **I)** Godet, “No”, “Yes” represent absence and presence of OB in the genotypes, respectively.

Similarly, there was variation for OB in the association panel at the two locations. OB occurs after the tuber is cut and exposed to oxygen (Figure 2A, 2B). A total of 46 and 38% of genotypes cultivated in Roujol and Godet showed OB, respectively. Godet might be a favorable environment for better-quality tubers.

**Figure 2.**
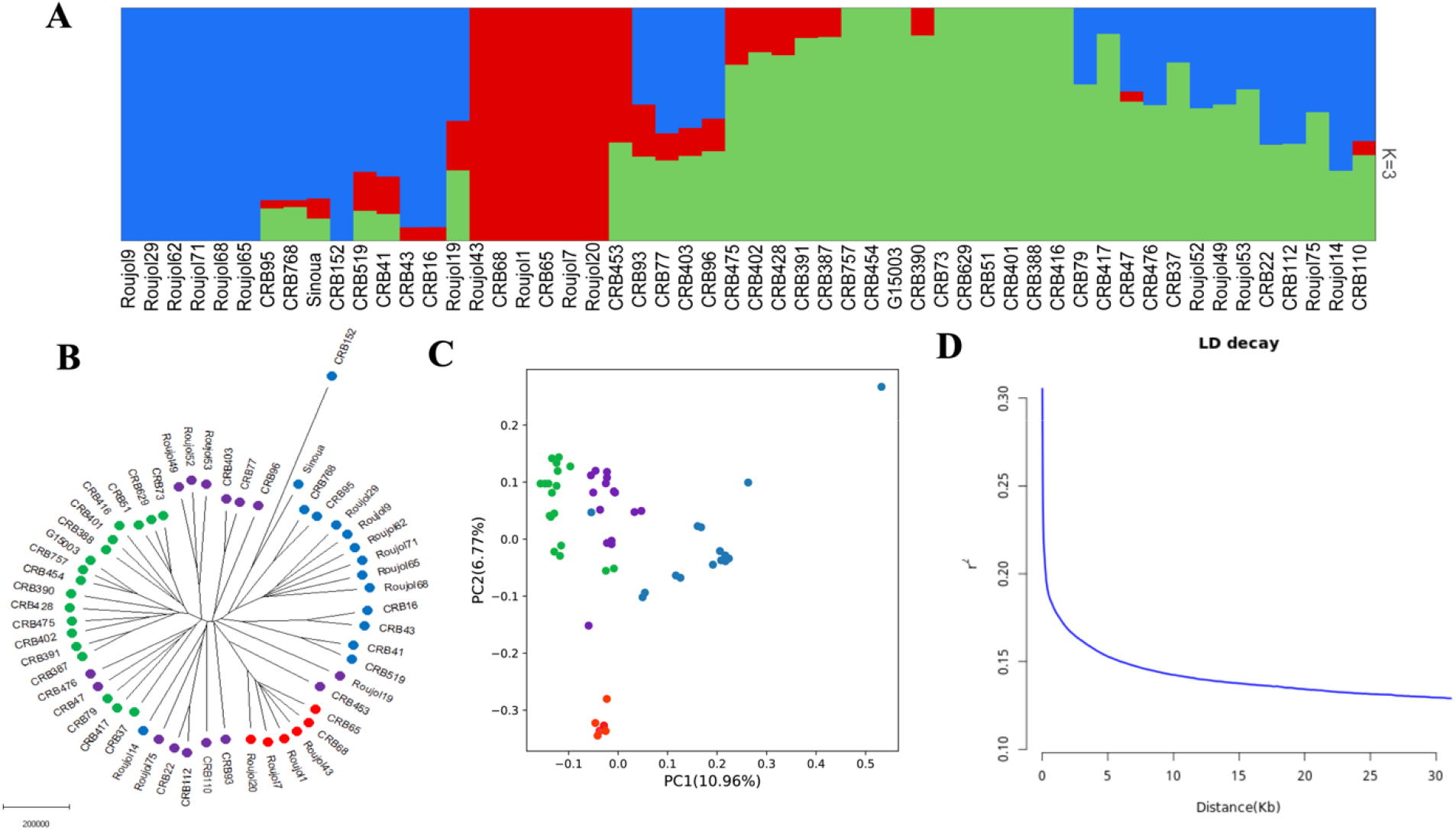
Population stratification analysis based on SNPs in *D. alata*. **A)** Model-based clustering analysis of all the genotypes performed using ADMIXTURE, representing three groups and their corresponding genotypes at K = 3, **B)** Circular dendrogram of the genotypes by the neighbor-joining method. *D. cayenensis* (CRB152) was used as an outgroup. Coloring followed the results from ADMIXTURE analysis, with purple-colored genotypes being the admixtures. The scale bar represents the length of the tree branch, **C)** Principal component analysis based on the high-quality SNPs. Coloring followed the results from ADMIXTURE analysis with purple-colored genotypes being the admixtures, **D)** LD decay estimates measured as the physical distance (kb) when LD decreased to half of its maximum value.

The Pearson correlation between both traits was very weak (r= -0.10), indicating that the tuber FC does not influence OB. Overall, we observed a significant variation among genotypes and environments, suggesting that the phenotypic data is suitable for GWAS.

### Genotyping data, population structure, and linkage disequilibrium

A total of 17 million single nucleotide polymorphisms (SNPs) were initially discovered between the genotypes of the association panel. After filtering, 1.9 million high-quality SNPs (minor allele frequency ≥ 0.05 and missing rate < 20%) were retained for genetic analysis, corresponding to a very high density of 3.9 SNPs/kb. The SNP distribution was relatively homogeneous along the chromosomes (Figure S1). However, as expected, the regions near centromeres have lower densities.

Population structure analysis showed the lowest cross-validation error at K=3 (Figure S2), indicating that the genotypes within the panel could be classified into 3 subpopulations, represented by the blue, red, and green colors (Figure 2A). Out of the 54 genotypes, only 21 pure genotypes were identified, with the remaining classified as admixed (Table S2). Overall, the diversity panel has a weak structure, which is favorable for genetic association analysis.

Similar to the population structure results, principal component analysis (PCA) and phylogenetic analysis did not show any obvious grouping of the studied genotypes (Figure 3C, 3D). The *D. cayenensis* (CRB152) used as an outgroup was clearly separated from the *D. alata* genotypes (Figure 2B). PCA showed weak clustering within the population, with the PC1 and PC2 covering only 17% of the total variation (Figure 2C). Linkage disequilibrium (LD) decay provides an estimate for the resolution of association mapping to assess the optimum numbers of SNPs. LD decay was observed at 5 kb (physical distance between SNPs) (Figure 2D).

**Figure 3.**
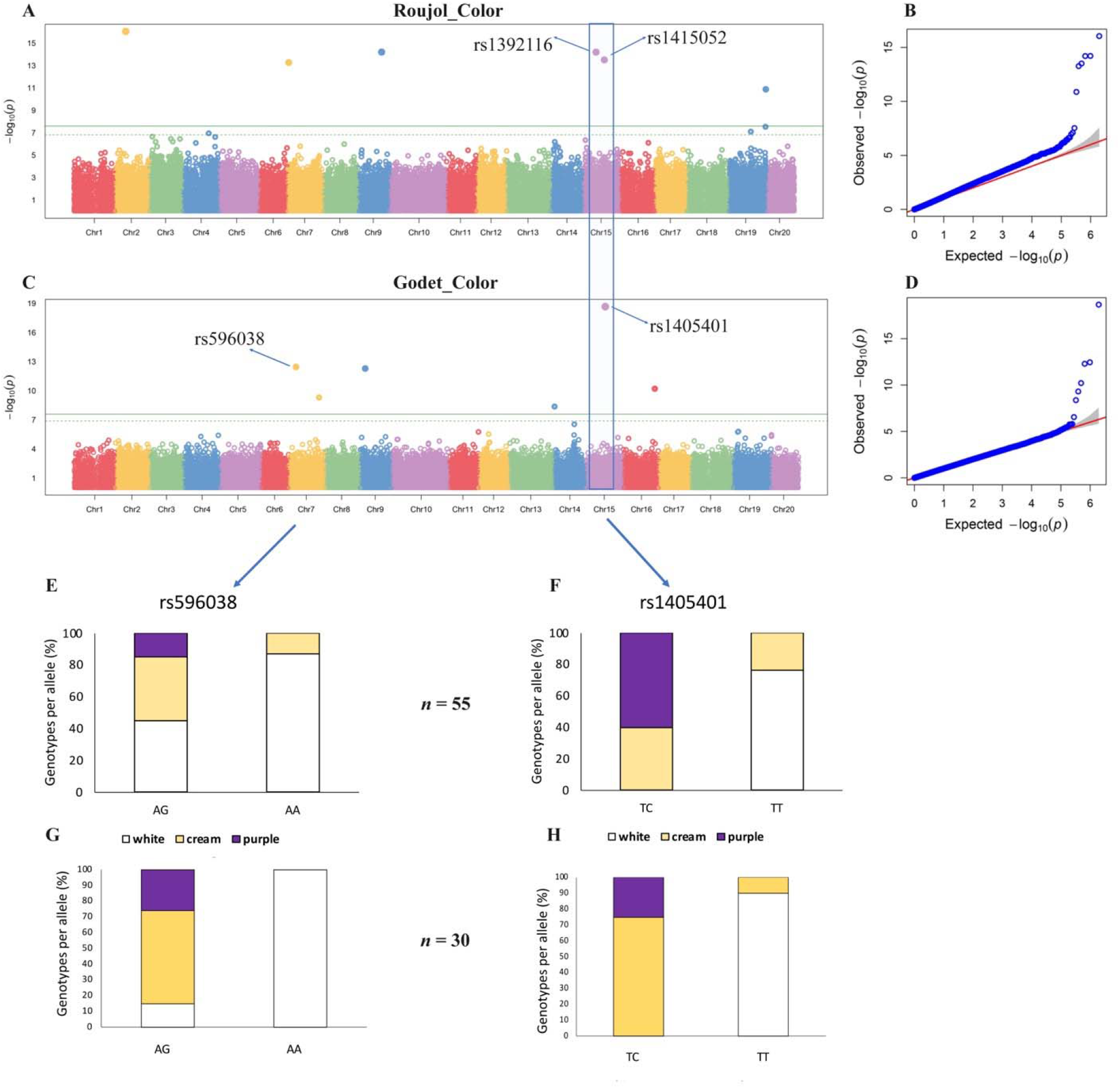
GWAS for tuber FC variation in *D. alata*. **A)** Manhattan plot for tuber FC at Roujol, with the peaks indicating significant GWAS signals, and the green horizontal lines indicating the genome-wide significance threshold, **B)** The QQ-Plot associated with tuber FC at Roujol shows the -log 10 of the expected vs. observed *P* values of each SNP (blue dots). The red line is a guide for the perfect fit to -log_10_*P*. The shaded area shows the 95% confidence interval for the QQ-plot under the null hypothesis of no association between the SNP and the trait, **C)** Manhattan plot for tuber FC at Godet, **D)** The QQ-plot associated with the tuber FC at Godet, **E & F)** Allele segregation analysis concerning SNPs rs596038 and rs1405401, where the x-axis represents the allelic variability at each locus and y-axis represents the proportion of genotypes representing the specific allele type. Purple, cream, and white colors indicate the corresponding genotypes with purple, cream, and white FCs, respectively, **G&H)** Allele segregation analysis concerning SNPs rs596038 and rs1405401 in an independent panel (size = 30 *D. alata* genotypes) grown and phenotyped in Vietnam.

Together, a large genetic diversity coupled with a low structure and a very weak LD decay were also observed in the association panel based on the SNP data (Figure 3), indicating that (1) the assembled panel is suitable for genome-wide association studies (GWAS) and (2) a high-density SNP is necessary to obtain reliable genotype-phenotype association results.

### Identification of loci controlling flesh color and oxidative browning in *D. alata*

The GWAS approach was implemented with the FarmCPU model to identify the genetic variants associated with the observed variation for FC and OB, with phenotypic data from each location independently.

### GWAS for tuber flesh color

We identified 14 significant associations (P< 10^−8^) for tuber FC using GWAS for the two locations (Table 1). At Roujol, seven significant associations were identified on Chr 2, Chr 7, Chr 9, Chr 15, and Chr 19 (Figure 3A, 4B), while at Godet, the significant associations were present on Chr 7, Chr 9, Chr 14, Chr 15, and Chr 16 (Figure 3C, 3D). Only, one genomic region on Chr 15 was commonly identified at the two locations. The discrepancy in the GWAS results (for the same trait and panel) at the two locations could be attributed to the environmental effects on the phenotype. To further understand the effect of these identified loci, we determined the allelic effect for the top significant or stable SNPs (rs596038 and rs1405401) using the allele segregation analysis.

**Table 1.** The significant association signals identified, with the allelic effects for each SNP.

| No | Site | Traits | SNP | CHR | Position | Allele | p-value (-log<sub>10</sub>) | Effect (%) |
| --- | --- | --- | --- | --- | --- | --- | --- | --- |
| 1 | Roujol | Flesh color | rs125884 | Chr2 | 7223285 | T/C | 16 | 17.40 |
| 2 | Roujol |  | rs832065 | Chr9 | 15404735 | G/GA | 14 | 13.95 |
| 3 | Roujol |  | rs1392116 | Chr15 | 8344629 | C/T | 14 | 9.69 |
| 4 | Roujol |  | rs1415052 | Chr15 | 13803586 | G/A | 14 | 8.41 |
| 5 | Roujol |  | rs575623 | Chr7 | 977139 | G/A | 13 | 8.87 |
| 6 | Roujol |  | rs1884180 | Chr19 | 24431097 | G/A | 11 | 8.24 |
| 7 | Roujol |  | rs1883574 | Chr19 | 24226102 | A/G | 75 | 7.69 |
| 8 | Godet |  | rs1405401 | Chr15 | 12347101 | T/TC | 18.67 | 18.04 |
| 9 | Godet |  | rs596038 | Chr7 | 4729592 | A/GA | 12.45 | 9.51 |
| 10 | Godet |  | rs766477 | Chr9 | 3312846 | T/C | 12.29 | 8.88 |
| 11 | Godet |  | rs1555677 | Chr16 | 20775954 | T/A | 10.21 | 7.42 |
| 12 | Godet |  | rs658251 | Chr7 | 19942330 | A/G | 9.31 | 7.18 |
| 13 | Godet |  | rs1275604 | Chr14 | 842033 | A/G | 8.37 | 7.67 |
| 14 | Roujol | Oxidative Browning | rs66743 | Chr1 | 21751133 | C/T | 12 | 12.27 |
| 15 | Roujol |  | rs1732730 | Chr18 | 20083173 | G/A | 11 | 11.53 |
| 16 | Roujol |  | rs176324<sup>£</sup> | Chr2 | 17011000 | G/T | 85 | 10.16 |
| 17 | Roujol |  | rs531469 | Chr6 | 8030455 | G/A | 77 | 7.75 |
| 18 | Godet |  | rs176324<sup>£</sup> | Chr2 | 17011000 | G/T | 7.21 | 10.76 |

**Figure 4.**
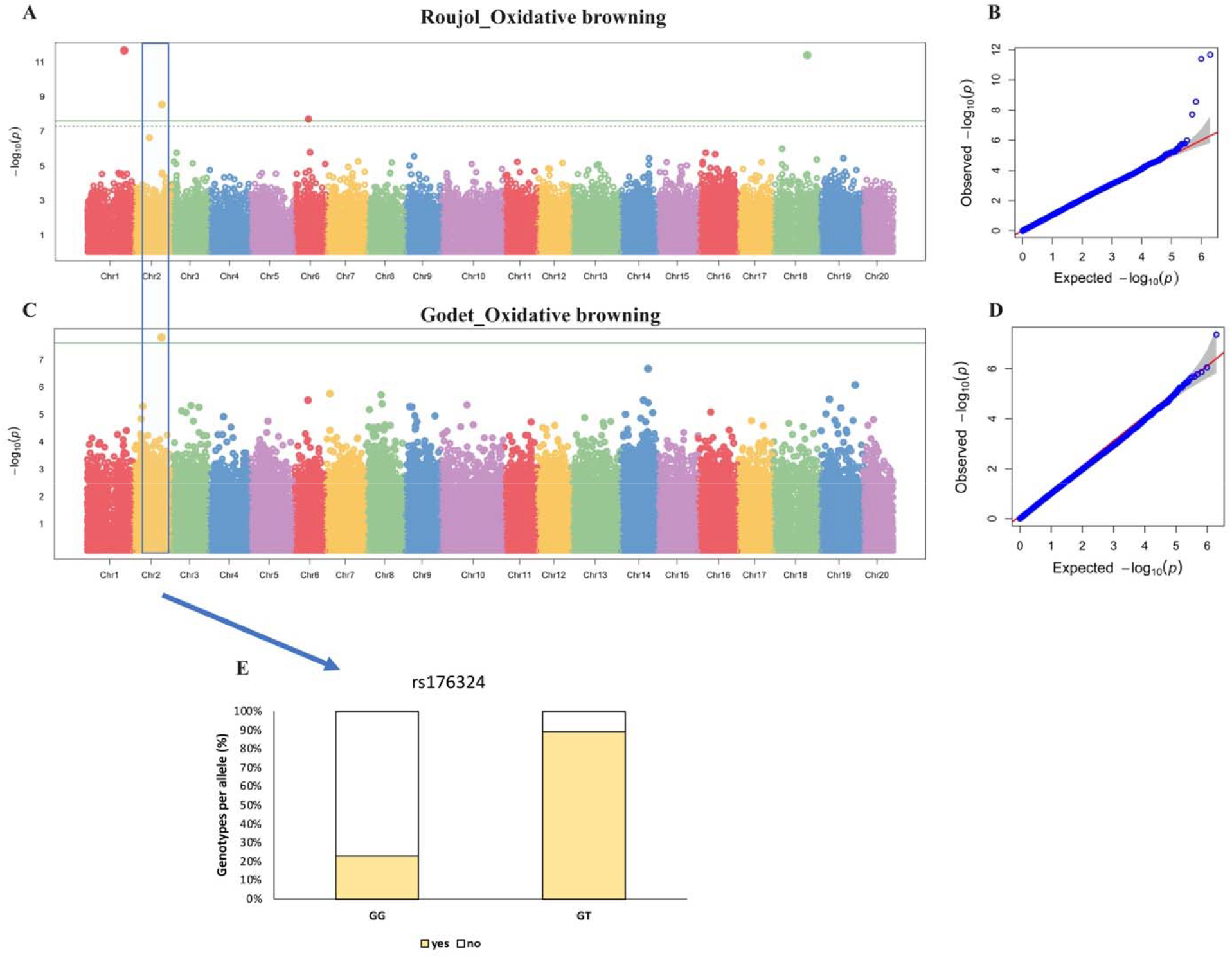
GWAS for OB in *D. alata*. **A) the** Manhattan plot for OB at Roujol, with the peaks indicating significant GWAS signals and the green horizontal lines indicating the genome-wide significance threshold, **B)** The QQ-plot associated with OB at Roujol, **C)** The Manhattan plot for OB at Godet, **D)** The QQ-plot plot associated with OB at Godet, **E)** Allele segregation analysis concerning SNP rs176324, where cream color indicates the genotypes with OB, and the white color indicates the genotypes without OB. “No”, “Yes” represent the absence and presence of OB in the genotypes, respectively.

### Allele segregation analysis for FC related peak SNPs

The two SNPs were consistently identified in close genomic regions at the two locations. The favorable alleles at each significant locus were identified, and the phenotypic variance explained was computed. For example, the detected significant SNP on chr 7 (rs596038) could explain 9.51% of the FC variation in our panel at Godet. We observed that the homozygous accessions (AA) at this locus have a significantly higher proportion of genotypes with white FC compared to the heterozygous accessions (AG) (Figure 3E). rs596038 is annotated as an intergenic SNP between the genes *Dioal*.*07G046300* (upstream) and *Dioal*.*07G046200* (downstream). *Dioal*.*07G046300* and *Dioal*.*07G046200* were annotated as *Calcium-binding protein 39* and *Auxin binding protein*, respectively (Table 2).

**Table 2.**
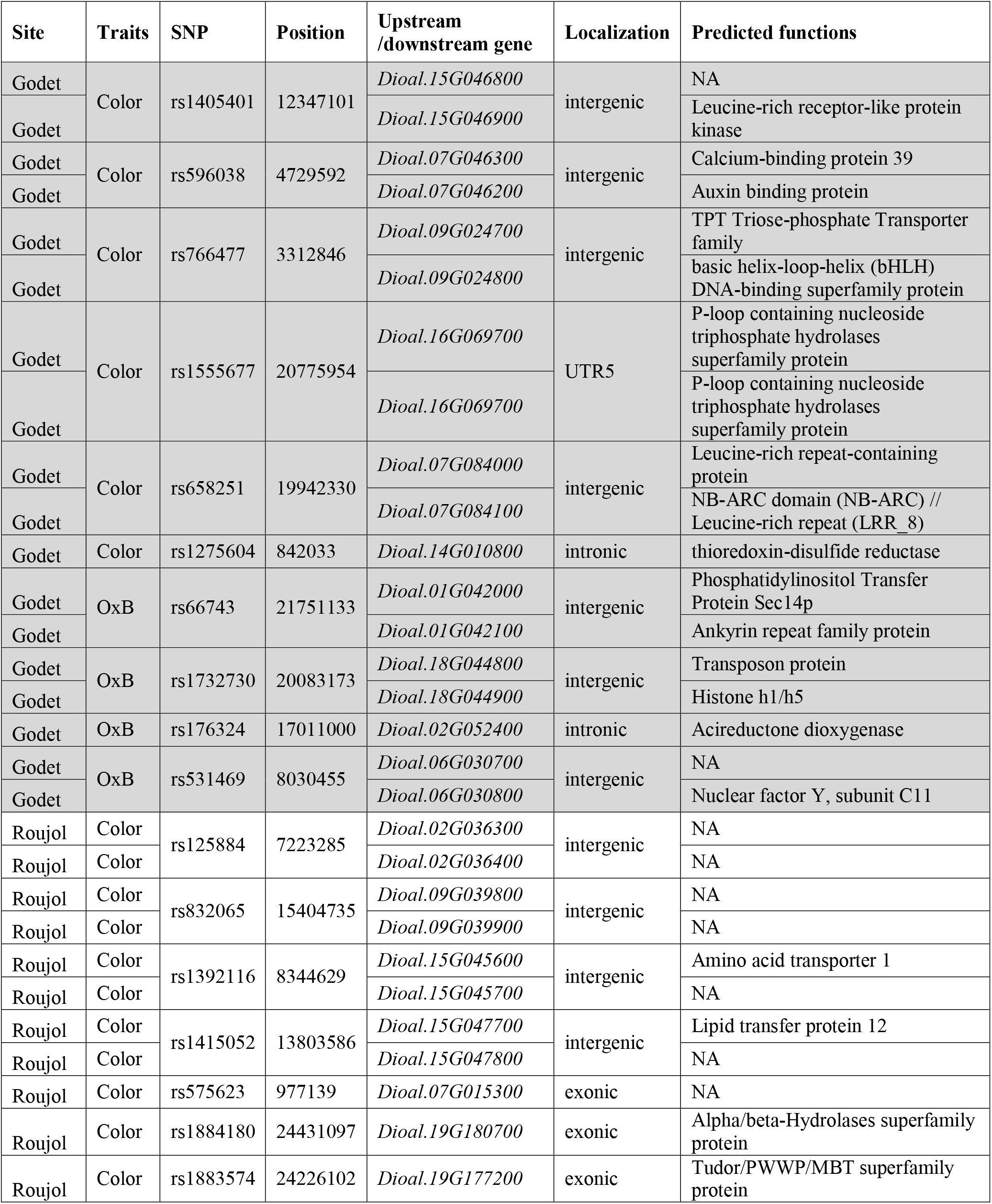
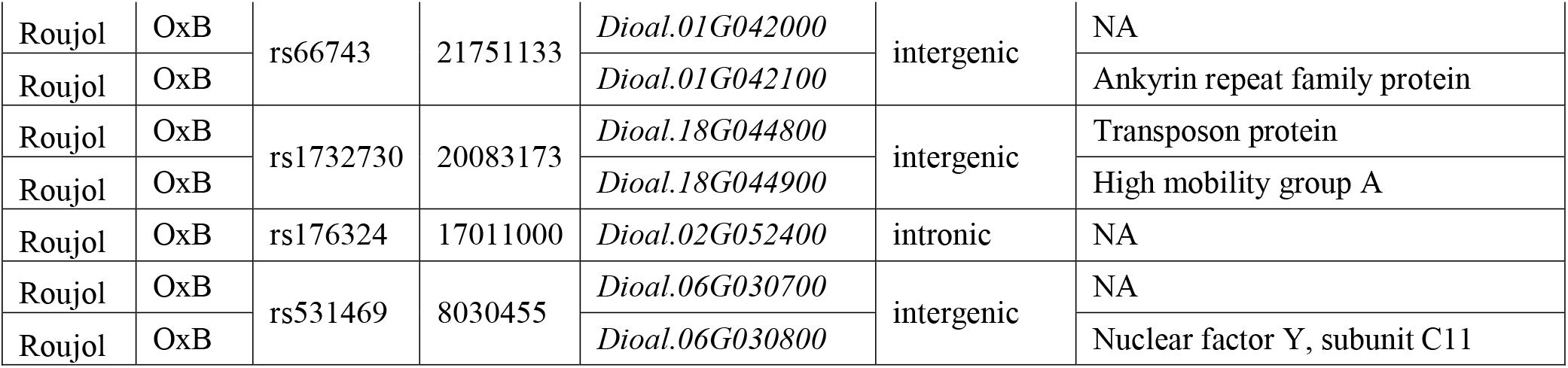
Summary of identified candidate genes associated with tuber FC and OB in *Dioscorea alata*.

Similarly, the significant SNP on chr 15 (rs1405401) was identified with TC and TT genotypes, where 80% of the accessions with homozygous TT genotype have white FC, while heterozygous TC genotype only corresponds to the accessions with cream and purple FCs. The phenotypic variance explained by rs1405401 was estimated to be 18.04% at Godet. Moreover, we identified around the SNP rs1405401 the *Dioal*.*15G046900* (annotated as *Leucine-rich receptor-like protein kinase*) and *Dioal*.*15G046800* (no annotation) in the downstream and upstream regions, respectively (Table 2). Allele segregation analysis for the remaining SNPs also depicted allele-specific phenotypes. For instance, SNPs rs832065, rs1392116, and rs1884180 were identified with specific alleles associated with the white tuber FC (Figure S3).

In order to validate the SNPs identified for FC and their allelic effect, we used an external panel of *D. alata* (*n* = 30) planted and phenotyped in Vietnam. The independent panel globally confirmed the allelic effect of identified GWAS loci on FC, suggesting that these loci could be used to predict FC in *D. alata* (Figure 3G and 3H).

### GWAS for oxidative browning

Four highly significant associations on Chr 1, Chr 2, Chr 6, and Chr 18 (Figures 4A, 4B, 4C, and 4D) for OB were found through GWAS in *D. alata*. To explore the genetic basis for tuber. Interestingly, the GWAS signal on Chr 2 (rs176324) was identified at both planting locations. The SNP effect of rs176324 was estimated to be 10.16% and 10.76% at Roujol and Godet, respectively (Table 1). We further explored the alleles of rs176324 and their effect on the tuber OB.

### Allele segregation analysis for OB-related peak SNPs

Allele segregation analysis suggested that *D. alata* genotypes with the homozygous loci GG does not display OB (Figure 5E). The allele “T” increases OB expression and, therefore, was considered unfavorable for breeding. The SNP rs176324 was identified in the intronic region of *Dioal*.*02G052400*, encoding *Acireductone dioxygenase* (Table 2). Allele segregation analysis for the remaining four SNPs also showed significant associations with OB; however, these SNPs were identified only at one location (Figure S4).

**Figure 5.**
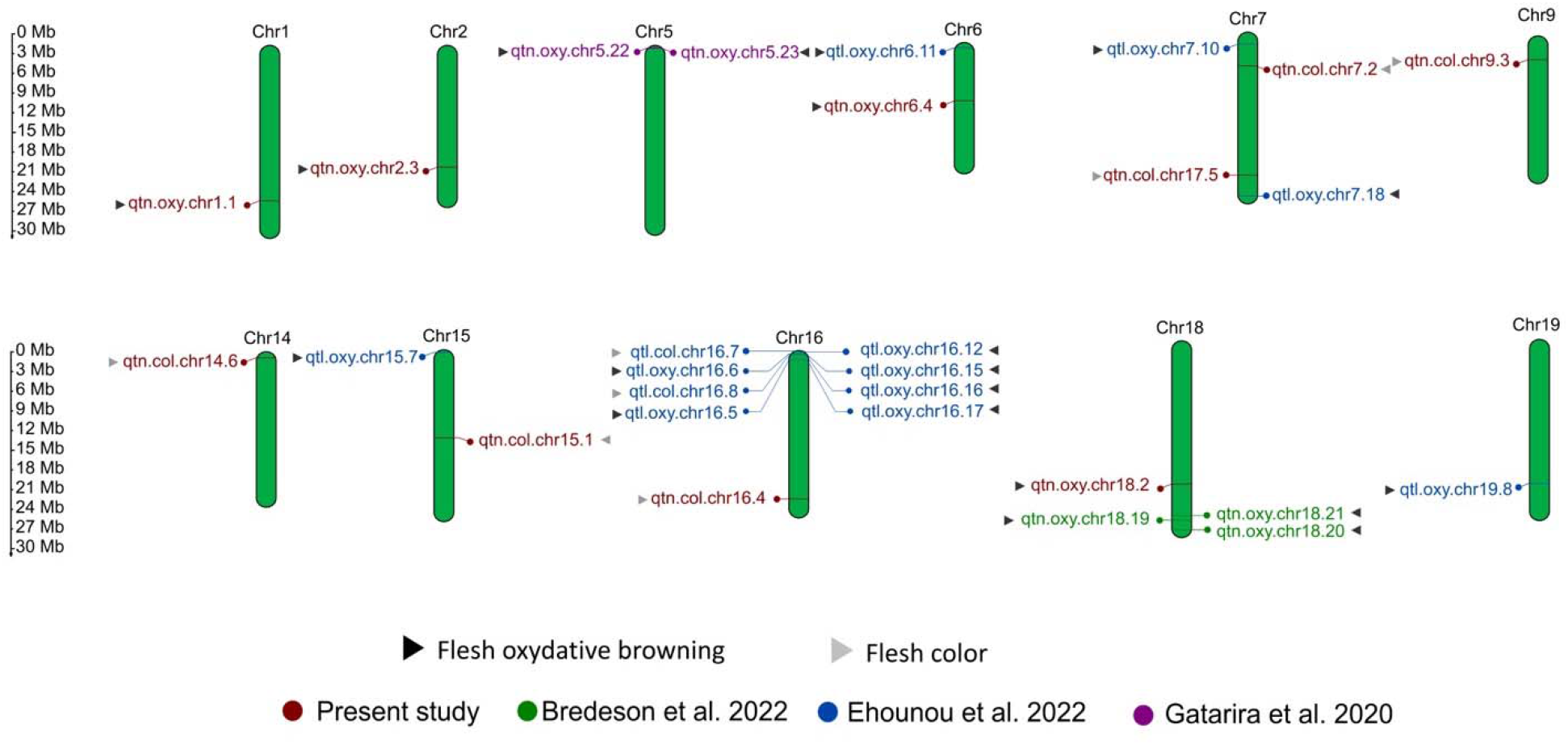
Genome-wide mapping (physical map) of quantitative trait loci from previous and current studies identified for tuber FC and OB in *D. alata* ^1,22,34^.

We also performed GWAS using the traditional Mixed Linear Model, however, the performance of FarmCPU was better in identifying the causal variants due to the highly heterozygous *D. alata* genome (Figure S5). Our results confirm a previous study where authors reported that FarmCPU is more robust than the traditional Mixed Linear Model in maize.^38^

### *In silico* comparative analysis of genomic regions identified from previous studies on tuber flesh color and oxidative browning

To further understand the extent of loci controlling the studied quality traits, we mapped the QTLs identified in the current study with those previously reported from three separate studies ^1,22,34^. Mapping results showed that none of the QTLs overlapped in the four studies. Although parental genotypes used by Bredeson et al., are present in the study of Gatarira et al., and similarly, parental genotypes in the study of Ehounou et al., are present in our panel, none of the QTLs coincides, emphasizing that FC and OB, are quantitative in nature, governed by several genomic regions and affected by genotype-by-environment interactions (Figure 5).

## Discussion

End-user quality should be a major objective in crop breeding programs.^39^ Unfortunately, there is a variable preference of FC in yam associated with regions and dishes, complicating breeding efforts considerably. For instance, white FC is the most preferred consumer trait in West Africa, while purple color is preferred in French Guyana, the Philippines, and India.^11,40^ Several studies have reported significant variations in the nutritional properties of different colored yams.^27,28^ Purple yam is enriched in anthocyanins, which are strong antioxidants providing additional nutritional value.^41^ As yam is widely consumed in West Africa, developing nutritionally enhanced cultivars would have a positive and significant impact on the diet of a large population. Therefore, increasing colorless anthocyanins in yam in order to maintain the desired white FC could be investigated. We observed an environmental influence on both traits, implying that the yam production environment is also important for obtaining optimal quality. Several studies have reported the environmental influence in managing yam yield^42,43^, however the environmental impact on yam quality traits are less documented. Identification of stable *D. alata* genotypes with desirable quality should be screened using multi-location trials for further use in breeding programs.

Recent GWAS reports in yams used panel sizes slightly higher than our study (74 and 80 accessions).^44,45^ Yams are diecious species with higher heterozygosity and allelic diversity as compared to hermaphrodite species. So even with relatively small sample sizes, it is possible to detect SNP-trait associations, especially with qualitative or not highly quantitative traits. In the diecious species Pistachio, only 44 accessions were used to successfully detect sex loci.^46^ In our initial analyses, simulation on different panel sizes to estimate the power for detecting true SNP-trait associations showed that when the variance explained by the locus was superior to 7%, all panels equally performed (data not shown). Hence, with a small panel, the only risk seems to not detect true SNP-trait associations for small effect loci. This is not problematic because small effect loci are not useful in marker-assisted breeding. Moreover, with a smaller panel, statistical power could be optimized by putting more efforts into accurate phenotyping. Sharif et al. demonstrated that from 643 worldwide *D. alata* accessions, only 90 accessions (14%) were non-clonally related.^2^ This means that having a large panel size dominated by true clones increases the population structure, biases estimation of allelic effects, and linkage disequilibrium (LD), which are crucial components for successful GWAS applications. Nonetheless, additional efforts are needed to enlarge our diversity panel with limited clonally related accessions to improve GWAS power and detect more marker-trait associations.

Based on the high-quality resequencing data, we observed a rapid LD-decay in *D. alata*. This is expected in dioecious species due to a high degree of recombination.^47^ Our results suggest that a high SNP density is required for association analyses. Therefore, we opted for a high SNP density (1.9 million SNPs, ∼3.9 SNPs/kb) in this study, which is significantly larger than any of the previous GWAS reports (less than 10,000 SNPs, ∼0.02 SNP/kb).^1^ High SNP-density with rapid LD-decay and a weak population structure have favored our GWAS.^48,49^ Thus, 18 significant associations with moderate effects (three of which were identified overlapped in both locations and could be considered stable QTLs) have been identified. The number of QTLs identified in this study combined with previous ones^1, 23,34^ and the evidence of a strong environmental influence, indicates that FC and OB are quantitative traits. These results are similar to those from Sun et al., for grape berry color.^50^ Allele segregation analysis has been extensively used to understand the effect of significant associations on population dynamics, providing insights to select advantageous alleles.^21,51^ In this study, allele segregation analysis at the significant loci further validated their association with the studied traits and highlighted the alleles to be considered in breeding programs.

A total of 24 genes around the significant GWAS signals, 14 for tuber FC and 10 for OB have been identified. *Dioal*.*07G046300, Dioal*.*07G046200*, and *Dioal*.*15G046900* encoding *Calcium-binding protein 39, Auxin binding protein*, and *Leucine-rich receptor-like protein kinase*, respectively, were identified as putative candidate genes for tuber FC. Pandey et al., demonstrated that *Calcium-binding protein* is involved in pigmentation in tobacco.^52^ Similarly, *Auxin binding protein* plays a significant role in plant tissue coloration.^53^ Interestingly, Hazak et al., discussed that *Calcium-binding protein* and its signaling pattern coincides with the expression pattern of Auxin-regulated genes.^54^ We hypothesize that *Dioal*.*07G046300* and *Dioal*.*07G046200*, located on the upstream and downstream regions of rs596038 on chromosome 15, might act harmoniously to regulate FC in *D. alata*. However, the underlying mechanisms are still not clear. Further functional study can elaborate on their potential roles in obtaining the desirable FC.

OB is a major quality issue in yam. Previous reports suggested that polyphenol oxidation (PPO) contributes to OB in yam.^16,18^ A recent study by Gatarira et al., identified several candidate genes associated with OB on chromosome 5.^1^ Contrary to their report, we did not find any significant association for OB on chromosome 5; instead, a stable signal on chromosome 2 was identified. Another study by Ehounou et al., used QTL analysis and identified several QTLs related to OB on chromosomes 5, 7, 15, 16, and 19.^22^ Bredeson et. al., identified three peroxidase-encoding genes (*Dioal*.*18G098800, Dioal*.*18G099400*, and *Dioal*.*18G100900*) on chromosome 18 using bi-parental mapping populations.^34^ The obvious differences between previous studies and our results are probably due to the different phenotyping procedures, genotyping methods and the quantitative nature of the trait. For example, two main phenotyping approaches (chromameter measurements^18^ and a visual scoring^22^) for yam tuber FC and OB have been used in the past. However, the correlation between these methods has not been investigated properly. We identified *Dioal*.*02G052400* encoding *Acireductone dioxygenase* as a putative candidate gene for tuber OB. *Acireductone dioxygenase* is a metal-binding metalloenzyme that binds with Fe^2+^ or Ni^2+^ in the methionine salvage pathway.^55^ Another report emphasized the role of *Acireductone dioxygenase* in ethylene biosynthesis.^56^ which is a key regulator for OB in crops through the modulation of PPO and other antioxidant enzymes.^57^ How ethylene potentially modulates polyphenol profiles leading to tuber OB in *D. alata*^18^ warrants further study.

Taken together, our study provided novel insights into the genetic control of FC and OB in *D. alata* tubers. Accessions accumulating favorable alleles at the different significant loci are valuable sources for future breeding programs. Nonetheless, our study showed that marker assisted breeding for these two quality traits could be difficult. This because numerous loci with moderate contributions are involved and there is a strong environmental influence. The potential of genomic selection on predicting FC and OB in *D. alata* should be investigated. Furthermore, we identified novel putative candidate genes for both traits. Functional analysis of these genes will contribute to a better understanding of their implications in FC and OB in *D. alata*.

### Editorial Policies and Ethical Considerations

All the experimental methods including field studies were performed in accordance with relevant guidelines and regulations.

## Supporting information

Density of SNPs along the 20 chromosomes of the Dioscorea alata genome. The horizontal axis indicates the length of the chromosomes, and the legend 0-

Plot of ADMIXTURE cross validation (CV) error from K=2 to K=10 in Dioscorea alata diversity panel. We selected K=3 as the value that minimizes the err

Allele segregation analysis for identified significant SNPs. A) rs125884, B) rs832065, C) rs 1392116, D) rs1883574, E) rs1884180, F) rs1415052, G) rs5

Allele segregation analysis for identified significant SNPs. A) rs66743, B) rs1732730, C) rs176324, D) rs531469, where the x-axis represents the allel

Manhattan plots of GWAS based on Mixed Linear Model for tuber FC and oxidative browning in Dioscorea alata at Roujol.

List of D. alata genotypes and D. cayenensis variety used in this study, along with their origins.

The proportion contributed by ancestral populations in each accession. A threshold of 60% was set to assign each accession to a Cluster.

## Acknowledgment

The authors are grateful to Marie-Claire Gravillon, Christophe Perrot and Elie Nudol for their contribution to the phenotyping. The editorial comments by Hernán Ceballos improved greatly the quality of this manuscript. This work was supported by the CGIAR Research Program on Roots, Tubers and Bananas (CRP-RTB) and the grant opportunity INV-008567 (formerly OPP1178942): Breeding RTB Products for End User Preferences (RTBfoods), to the French Agricultural Research Centre for International Development (CIRAD), Montpellier, France, by the Bill & Melinda Gates Foundation (BMGF): https://rtbfoods.cirad.fr.

## Conflicts of Interest

The authors declare no competing financial interests.

## Data availability

The Illumina NovaSeq 6000 sequencing raw data are available in the NCBI Sequence Read Archive, under the BioProject number: PRJNA880983. Data will be released upon publication of this manuscript. The phenotypic datasets are available from the corresponding author upon request.

## Supplementary materials

**Figure S1**. Density of SNPs along the 20 chromosomes of the *Dioscorea alata* genome. The horizontal axis indicates the length of the chromosomes, and the legend 0-8865 indicates the density of the SNPs.

**Figure S2**. Plot of ADMIXTURE cross validation (CV) error from K=2 to K=10 in *Dioscorea alata* diversity panel. We selected K=3 as the value that minimizes the error.

**Figure S3**. Allele segregation analysis for identified significant SNPs. **A**) rs125884, **B**) rs832065, **C**) rs 1392116, **D**) rs1883574, **E**) rs1884180, **F**) rs1415052, **G**) rs575623, where the x-axis represents the allelic variability at the locus, and the y-axis represents the proportion of *Dioscorea alata* genotypes representing the specific allele type. Purple, cream, and white colors indicate the corresponding genotypes with purple, cream and white FCs, respectively.

**Figure S4**. Allele segregation analysis for identified significant SNPs. **A**) rs66743, **B**) rs1732730, **C**) rs176324, **D**) rs531469, where the x-axis represents the allelic variability at the locus, and the y-axis represents the proportion of genotypes representing the specific allele type. Cream color indicates the genotypes with oxidative browning, and the white color indicates the genotypes without oxidative browning.

**Figure S5**. Manhattan plots of GWAS based on Mixed Linear Model for tuber FC and oxidative browning in *Dioscorea alata* at Roujol.

**Table S1**. List of *D. alata* genotypes and *D. cayenensis* variety used in this study, along with their origins.

**Table S2**. The proportion contributed by ancestral populations in each accession. A threshold of 60% was set to assign each accession to a Cluster.

## Notes

### Competing Interest Statement

The authors have declared no competing interest.

## References

1. Gatarira C, Agre P, Matsumoto R, Edemodu A, Adetimirin V, Bhattacharjee R, Asiedu R and Asfaw A, Genome-wide association analysis for tuber dry matter and oxidative browning in water yam (Dioscorea alata L.). Plants 9:969 (2020).

2. Sharif BM, Burgarella C, Cormier F, Mournet P, Causse S, Van KN, Kaoh J, Rajaonah MT, Lakshan SR and Waki J, Genome-wide genotyping elucidates the geographical diversification and dispersal of the polyploid and clonally propagated yam (Dioscorea alata). Annals of botany 126:1029–1038 (2020).

3. Srivastava AK, Gaiser T, Paeth H and Ewert F, The impact of climate change on Yam (Dioscorea alata) yield in the savanna zone of West Africa. Agriculture, ecosystems & environment 153:57–64 (2012).

4. Pouya N, Hgaza VK, Kiba DI, Bomisso L, Aighewi B, Aké S and Frossard E, Water yam (Dioscorea alata L.) growth and tuber yield as affected by rotation and fertilization regimes across an environmental gradient in west Africa. Agronomy 12:792 (2022).

5. Cornet D, Marcos J, Tournebize R and Sierra J, Observed and modeled response of water yam (Dioscorea alata L.) to nitrogen supply: Consequences for nitrogen fertilizer management in the humid tropics. European Journal of Agronomy 138:126536 (2022).

6. Otoo E, Opoku-Agyeman M, Dansi A, Aboagye L, Acheremu K and Tetteh J, Increasing farmers and breeders access to yam (Dioscorea Spp) Diversity: The case of forest-savannah transition agroecology. African Journal of Agricultural Research 10:772–782 (2015).

7. Egesi CN, Asiedu R, Egunjobi JK and Bokanga M, Genetic diversity of organoleptic properties in water yam (Dioscorea alata L). Journal of the Science of Food and Agriculture 83:858–865 (2003).

8. Effah-Manu L, Wireko-Manu FD, Agbenorhevi JK, Maziya-Dixon B, Oduro IN and Baah-Ennum TY, The effect of gender on end-user preferences for yam quality descriptors. Food Science & Nutrition (2022).

9. Clawson DL, Harvest security and intraspecific diversity in traditional tropical agriculture. Economic Botany 39:56–67 (1985).

10. Champagne A, Legendre L and Lebot V, Chemotype profiling to guide breeders and explore traditional selection of tropical root crops in Vanuatu, South Pacific. Journal of agricultural and food chemistry 57:10363–10370 (2009).

11. Lebot V, Tropical root and tuber crops. Cabi (2019).

12. Wang A, Li R, Ren L, Gao X, Zhang Y, Ma Z, Ma D and Luo Y, A comparative metabolomics study of flavonoids in sweet potato with different flesh colors (Ipomoea batatas (L.) Lam). Food chemistry 260:124–134 (2018).

13. Tanaka Y, Sasaki N and Ohmiya A, Biosynthesis of plant pigments: anthocyanins, betalains and carotenoids. The Plant Journal 54:733–749 (2008).

14. Lebot V, Malapa R, Molisalé T and Marchand J-L, Physico-chemical characterisation of yam (Dioscorea alata L.) tubers from Vanuatu. Genetic Resources and Crop Evolution 53:1199–1208 (2006).

15. Shittu TA and Olaitan OF, Functional effects of dried okra powder on reconstituted dried yam flake and sensory properties of ojojo—a fried yam (Dioscorea alata L.) snack. Journal of food science and technology 51:359–364 (2014).

16. Martin FW and Ruberte R, Polyphenol of Dioscorea alata (yam) tubers associated with oxidative browning. Journal of Agricultural and food Chemistry 24:67–70 (1976).

17. Srivichai S and Hongsprabhas P, Profiling anthocyanins in Thai purple yams (Dioscorea alata L.). International Journal of Food Science 2020 (2020).

18. Rinaldo D, Sotin H, Pétro D, Le-Bail G and Guyot S, Browning susceptibility of new hybrids of yam (Dioscorea alata) as related to their total phenolic content and their phenolic profile determined using LC-UV-MS. LWT 162:113410 (2022).

19. Ray DK, Ramankutty N, Mueller ND, West PC and Foley JA, Recent patterns of crop yield growth and stagnation. Nature communications 3:1–7 (2012).

20. Voss-Fels KP, Herzog E, Dreisigacker S, Sukumaran S, Watson A, Frisch M, Hayes B and Hickey LT, “SpeedGS” to accelerate genetic gain in spring wheat, in Applications of genetic and genomic research in cereals. Elsevier, pp 303–327 (2019).

21. Nazir MF, He S, Ahmed H, Sarfraz Z, Jia Y, Li H, Sun G, Iqbal MS, Pan Z and Du X, Genomic insight into the divergence and adaptive potential of a forgotten landrace G. áhirsutum L. purpurascens. Journal of Genetics and Genomics 48:473–484 (2021).

22. Ehounou AE, Cormier F, Maledon E, Nudol E, Vignes H, Gravillon MC, N’guetta Asp, Mournet P, Chaïr H and Kouakou AM, Identification and validation of QTLs for tuber quality related traits in greater yam (Dioscorea alata L.). Scientific Reports 12:1–14 (2022).

23. Andrews S, FastQC: a quality control tool for high throughput sequence data, Ed. Babraham Bioinformatics, Babraham Institute, Cambridge, United Kingdom (2010).

24. Bolger AM, Lohse M and Usadel B, Trimmomatic: a flexible trimmer for Illumina sequence data. Bioinformatics 30:2114–2120 (2014).

25. Li H, Exploring single-sample SNP and INDEL calling with whole-genome de novo assembly. Bioinformatics 28:1838–1844 (2012).

26. Okonechnikov K, Conesa A and García-Alcalde F, Qualimap 2: advanced multi-sample quality control for high-throughput sequencing data. Bioinformatics 32:292–294 (2016).

27. McKenna A, Hanna M, Banks E, Sivachenko A, Cibulskis K, Kernytsky A, Garimella K, Altshuler D, Gabriel S and Daly M, The Genome Analysis Toolkit: a MapReduce framework for analyzing next-generation DNA sequencing data. Genome research 20:1297–1303 (2010).

28. Danecek P, Auton A, Abecasis G, Albers CA, Banks E, DePristo MA, Handsaker RE, Lunter G, Marth GT and Sherry ST, The variant call format and VCFtools. Bioinformatics 27:2156–2158 (2011).

29. Yin L, CMplot: circle manhattan plot. R package version 3 (2020).

30. Team RC, R Foundation for Statistical Computing; Vienna: 2018. R: a language and environment for statistical computing[Google Scholar] (2016).

31. Bender D, Maller J, Sklar P, de Bakker P, Daly M and Sham P, PLINK: a toolset for whole-genome association and population-based linkage analysis. Am J Hum Genet 81:559–575 (2007).

32. Alexander DH, Novembre J and Lange K, Fast model-based estimation of ancestry in unrelated individuals. Genome research 19:1655–1664 (2009).

33. Wang K, Li M and Hakonarson H, ANNOVAR: Functional annotation of genetic variants from next-generation sequencing data Nucleic Acids Research. ANNOVAR: Functional annotation of genetic variants from next-generation sequencing data Nucleic Acids Research 38 (2010).

34. Bredeson JV, Lyons JB, Oniyinde IO, Okereke NR, Kolade O, Nnabue I, Nwadili CO, Hřibová E, Parker M and Nwogha J, Chromosome evolution and the genetic basis of agronomically important traits in greater yam. Nature communications 13:1–16 (2022).

35. Wang J and Zhang Z, GAPIT version 3: boosting power and accuracy for genomic association and prediction. Genomics, proteomics & bioinformatics 19:629–640 (2021).

36. Liu X, Huang M, Fan B, Buckler ES and Zhang Z, Iterative usage of fixed and random effect models for powerful and efficient genome-wide association studies. PLoS genetics 12:e1005767 (2016).

37. Voorrips R, MapChart: software for the graphical presentation of linkage maps and QTLs. Journal of heredity 93:77–78 (2002).

38. Cui Z, Dong H, Zhang A, Ruan Y, Jiang S, He Y and Zhang Z, Denser markers and advanced statistical method identified more genetic loci associated with husk traits in maize. Scientific reports 10:1–10 (2020).

39. Randhawa HS, Bona L and Graf R, Triticale breeding—Progress and prospect, in Triticale. Springer, pp 15–32 (2015).

40. Otegbayo B, Madu T, Oroniran O, Chijioke U, Fawehinmi O, Okoye B, Tanimola A, Adebola P and Obidiegwu J, End-user preferences for pounded yam and implications for food product profile development. International Journal of Food Science & Technology 56:1458–1472 (2021).

41. Chen X, Sun J, Zhu Q, Xiao Y, Zhang H, Huang Y, Wang P, Cao T, Hu R and Xiang Z, Characterizing diversity based on phenotypes and molecular marker analyses of purple yam (Dioscorea alata L.) germplasm in southern China. Genetic Resources and Crop Evolution:1–13 (2022).

42. Asfaw A, Standard operating protocol for yam variety performance evaluation trial, Ed (2016).

43. Norman PE, Tongoona PB, Danquah A, Danquah EY, Agre PA, Agbona A, Asiedu R and Asfaw A, Genetic Analysis of Agronomic and Quality Traits from Multi-Location white Yam Trials using Mixed Model with Genomic Relationship Matrix. Global Journal of Botanical Science 10:8–22 (2022).

44. Mondo JM, Agre PA, Asiedu R, Akoroda MO and Asfaw A, Genome-wide association studies for sex determination and cross-compatibility in water yam (Dioscorea alata L.). Plants 10:1412 (2021).

45. Cormier, F, Lawac, F, Maledon, E, Gravillon, MC, Nudol, E, Mournet, P, Vignes, H, Chaïr, H, and Arnau, G, A reference high-density genetic map of greater yam (Dioscorea alata L.). Theor Appl Genet 132: 1733–1744 (2019).

46. Kafkas S, Ma X, Zhang X, Topçu H, Navajas-Pérez R, Wai CM, Tang H, Xu X, Khodaeiaminjan M and Güney M, Pistachio genomes provide insights into nut tree domestication and ZW sex chromosome evolution. Plant Communications:100497 (2022).

47. Barnaud A, Laucou V, This P, Lacombe T and Doligez A, Linkage disequilibrium in wild French grapevine, Vitis vinifera L. subsp. silvestris. Heredity 104:431–437 (2010).

48. Migicovsky Z, Gardner KM, Money D, Sawler J, Bloom JS, Moffett P, Chao CT, Schwaninger H, Fazio G and Zhong GY, Genome to phenome mapping in apple using historical data. The Plant Genome 9:plantgenome2015.2011.0113 (2016).

49. Sul JH, Martin LS and Eskin E, Population structure in genetic studies: Confounding factors and mixed models. PLoS genetics 14:e1007309 (2018).

50. Sun L, Li S, Jiang J, Tang X, Fan X, Zhang Y, Liu J and Liu C, New quantitative trait locus (QTLs) and candidate genes associated with the grape berry color trait identified based on a high-density genetic map. BMC Plant Biology 20:1–13 (2020).

51. Liu Y, Nazir MF, He S, Li H, Pan Z, Sun G, Dai P, Wang L and Du X, Deltapine 15 contributes to the genomic architecture of modern upland cotton cultivars. Theoretical and Applied Genetics 135:1401–1411 (2022).

52. Pandey GK, Pandey A, Reddy VS, Deswal R, Bhattacharya A, Upadhyaya KC and Sopory SK, Antisense expression of a gene encoding a calcium-binding protein in transgenic tobacco leads to altered morphology and enhanced chlorophyll. Journal of biosciences 32:251–260 (2007).

53. Tan L, Tan C, Zuraida A, Hossein H, Goh H, Ismanizan I and Zamri Z, Overexpression of auxin binding protein 57 from Rice (Oryza sativa L.) increased drought and salt tolerance in transgenic Arabidopsis thaliana, in IOP Conference Series: Earth and Environmental Science, Ed. IOP Publishing, p 012038 (2018).

54. Hazak O, Mamon E, Lavy M, Sternberg H, Behera S, Schmitz-Thom I, Bloch D, Dementiev O, Gutman I and Danziger T, A novel Ca2+-binding protein that can rapidly transduce auxin responses during root growth. PLoS biology 17:e3000085 (2019).

55. Liang S, Xiong W, Yin C, Xie X, Jin Y-j, Zhang S, Yang B, Ye G, Chen S and Luan W-j, Overexpression of OsARD1 improves submergence, drought, and salt tolerances of seedling through the enhancement of ethylene synthesis in rice. Frontiers in plant science 10:1088 (2019).

56. Guo T, Zhang X, Li Y, Liu C, Wang N, Jiang Q, Wu J, Ma F and Liu C, Overexpression of MdARD4 Accelerates Fruit Ripening and Increases Cold Hardiness in Tomato. International journal of molecular sciences 21:6182 (2020).

57. Jung S-K and Watkins CB, Involvement of ethylene in browning development of controlled atmosphere-stored ‘Empire’apple fruit. Postharvest biology and technology 59:219–226 (2011).

