## Supplementary figures and images for "Genome-wide association studies reveal novel loci controlling tuber flesh color and oxidative browning in *Dioscorea alata*"

### Allele segregation analysis for identified significant SNPs. A) rs66743, B) rs1732730, C) rs176324, D) rs531469, where the x-axis represents the allel

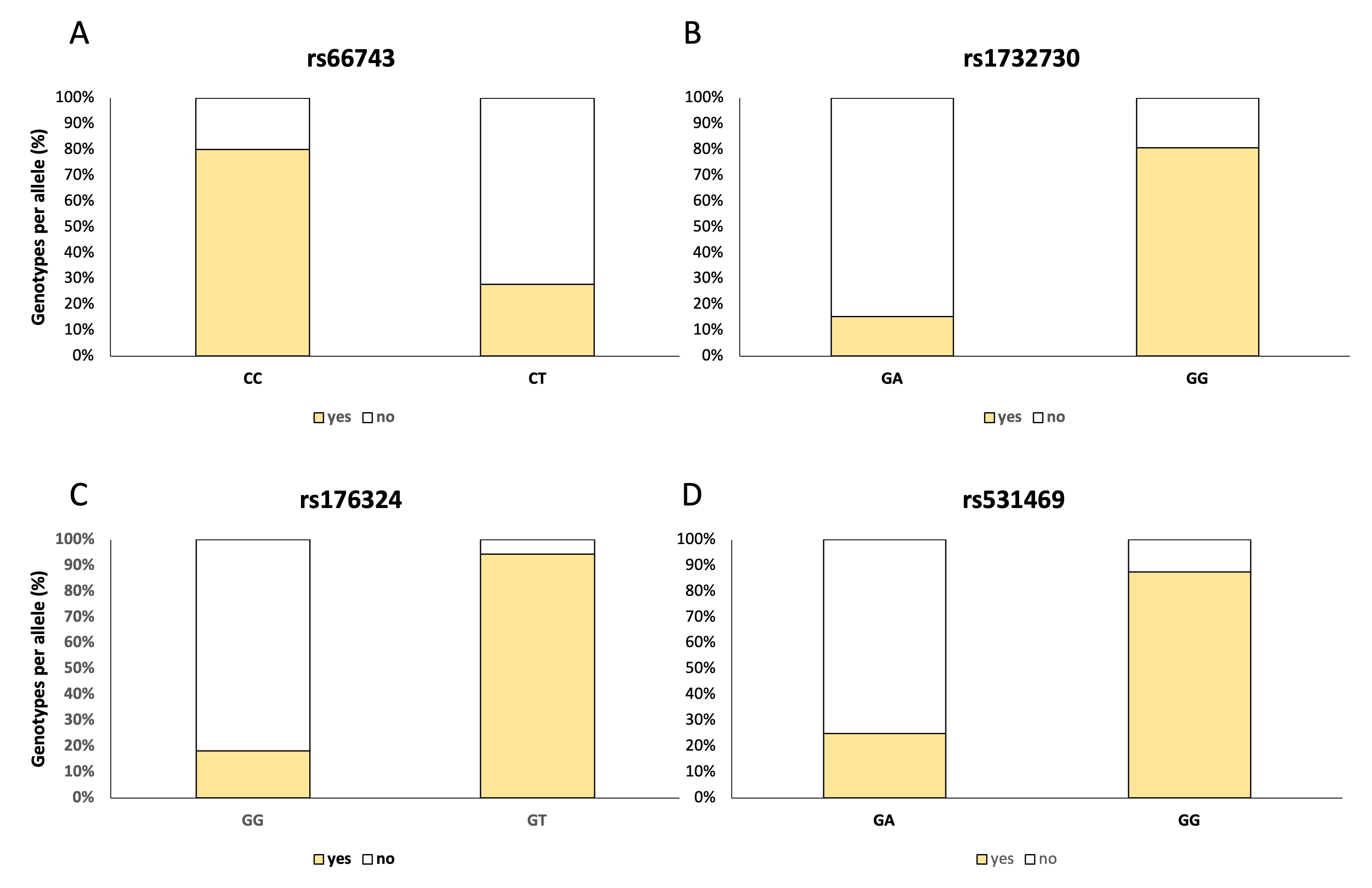

### Allele segregation analysis for identified significant SNPs. A) rs125884, B) rs832065, C) rs 1392116, D) rs1883574, E) rs1884180, F) rs1415052, G) rs5

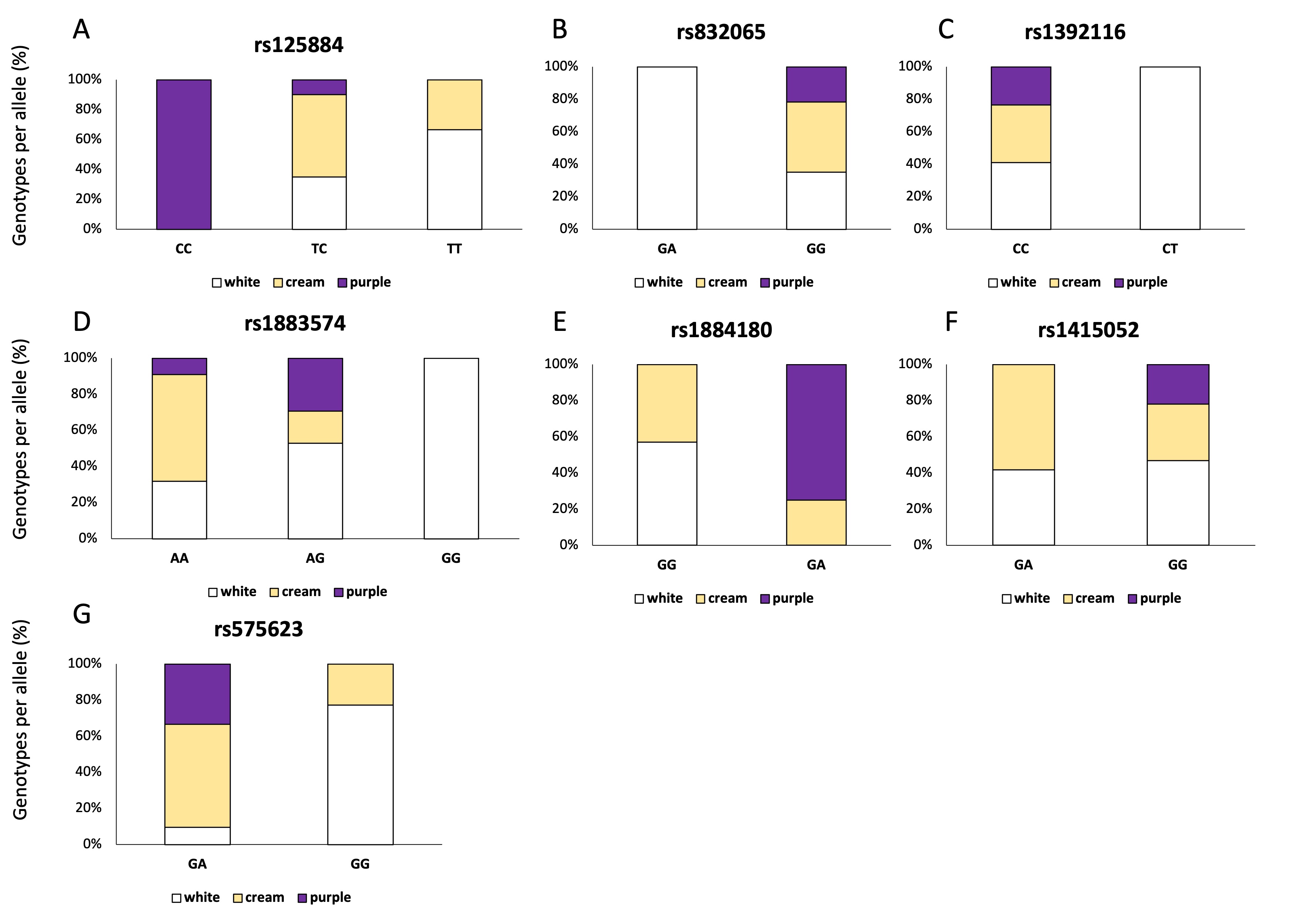

### Density of SNPs along the 20 chromosomes of the Dioscorea alata genome. The horizontal axis indicates the length of the chromosomes, and the legend 0-

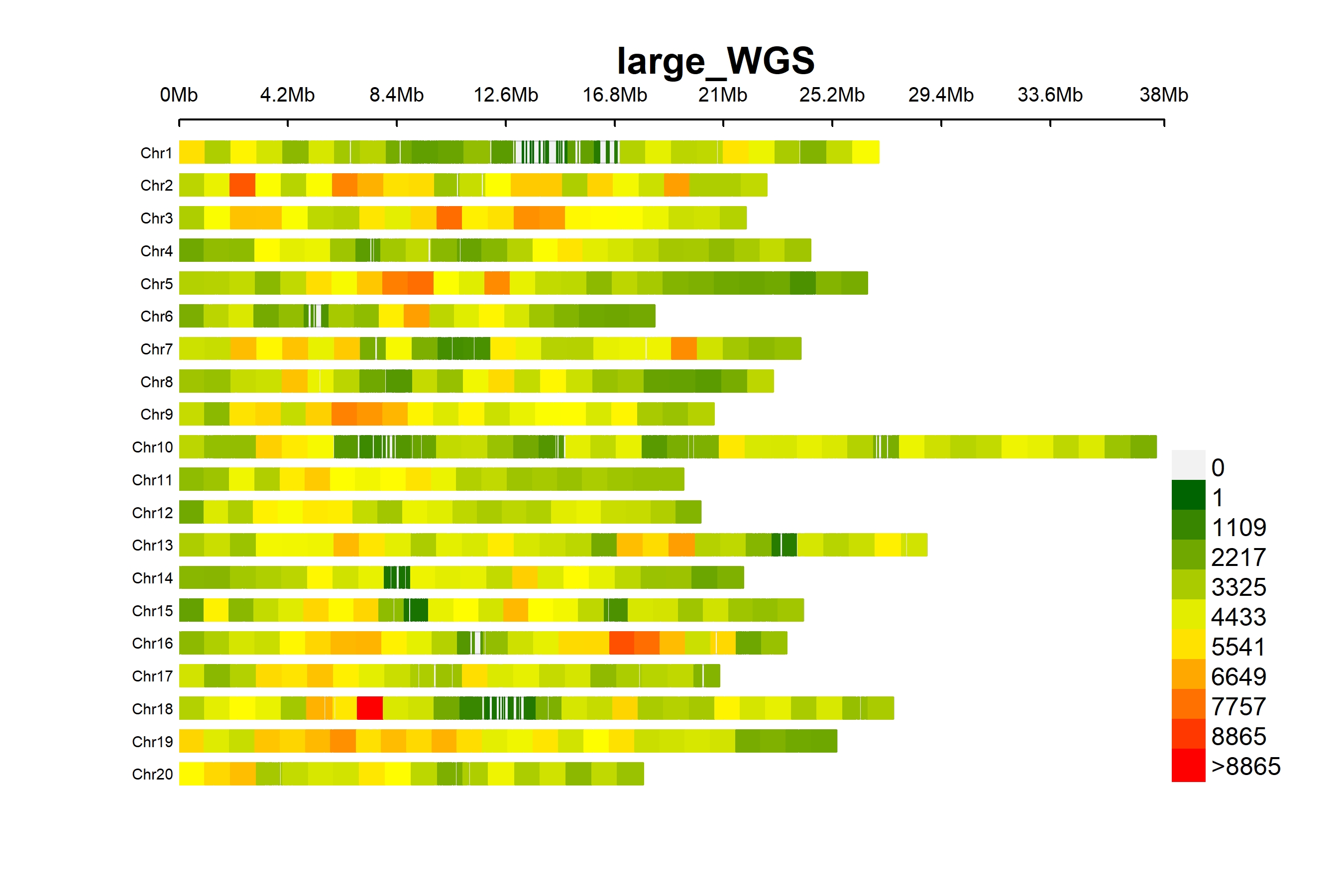

### Manhattan plots of GWAS based on Mixed Linear Model for tuber FC and oxidative browning in Dioscorea alata at Roujol.

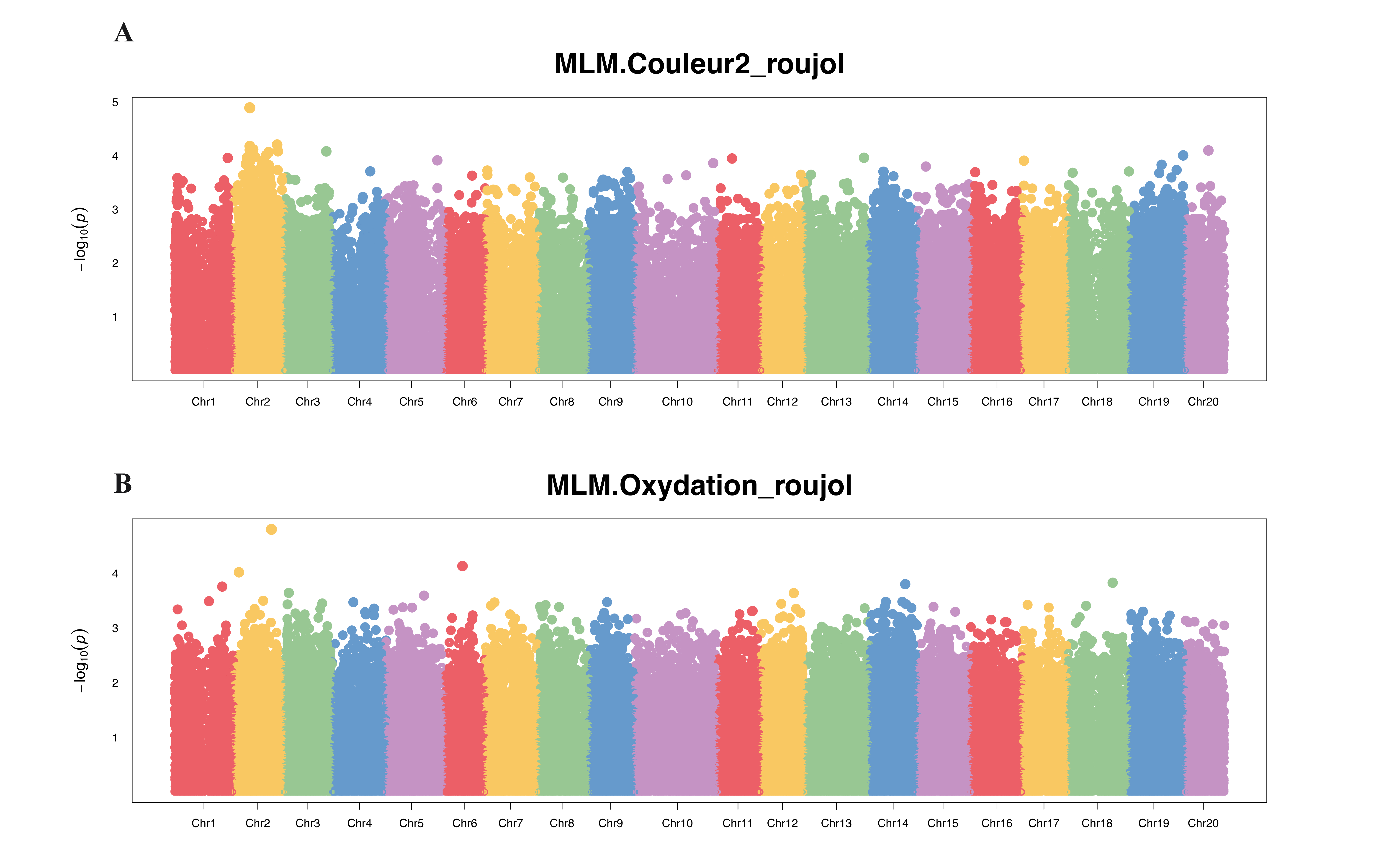

### Plot of ADMIXTURE cross validation (CV) error from K=2 to K=10 in Dioscorea alata diversity panel. We selected K=3 as the value that minimizes the err

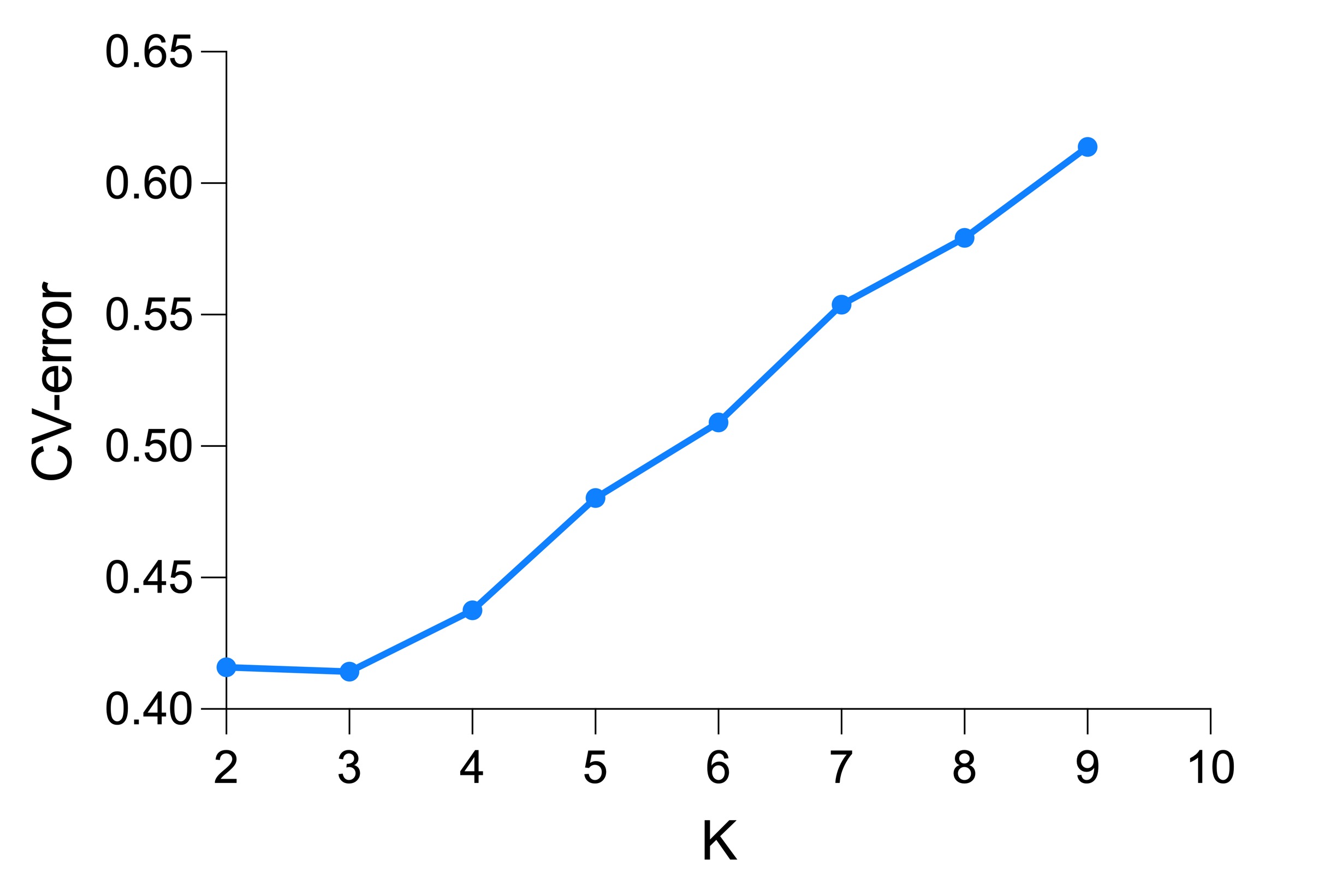
